# Amorphous solid dispersions and the confounding effect of nanoparticles in *in vitro* dissolution and in vivo testing: Niclosamide as a case study

**DOI:** 10.1101/2020.12.22.424004

**Authors:** Miguel O. Jara, Zachary N. Warnken, Robert O. Williams

## Abstract

We developed an amorphous solid dispersion (ASD) of the poorly water-soluble molecule niclosamide that achieved more than a 2-fold increase in bioavailability. Notably, this niclosamide ASD formulation increased the apparent drug solubility about 60-fold relative to the crystalline material due to the generation of nanoparticles. Niclosamide is a weakly acidic drug, BCS class II, and a poor glass former with low bioavailability in vivo. Hot-melt extrusion is a high-throughput manufacturing method commonly used in the development of ASDs for increasing the apparent solubility and bioavailability of poorly water-soluble compounds. We utilized the polymer polyvinylpyrrolidone–vinyl acetate (PVP–VA) to manufacture niclosamide ASDs by extrusion. Samples were analyzed based on their microscopic and macroscopic behavior and their intermolecular interactions, using DSC, XRD, NMR, FTIR, and DLS. The niclosamide ASD generated nanoparticles with a mean particle size of about 100 nm in FaSSIF media. In a side-by-side diffusion test, these nanoparticles produced a 4-fold increase in niclosamide diffusion. We successfully manufactured amorphous extrudates of the poor glass former niclosamide that showed remarkable in vitro dissolution and diffusion performance. These in vitro tests were translated to a rat model that also showed an increase in oral bioavailability.

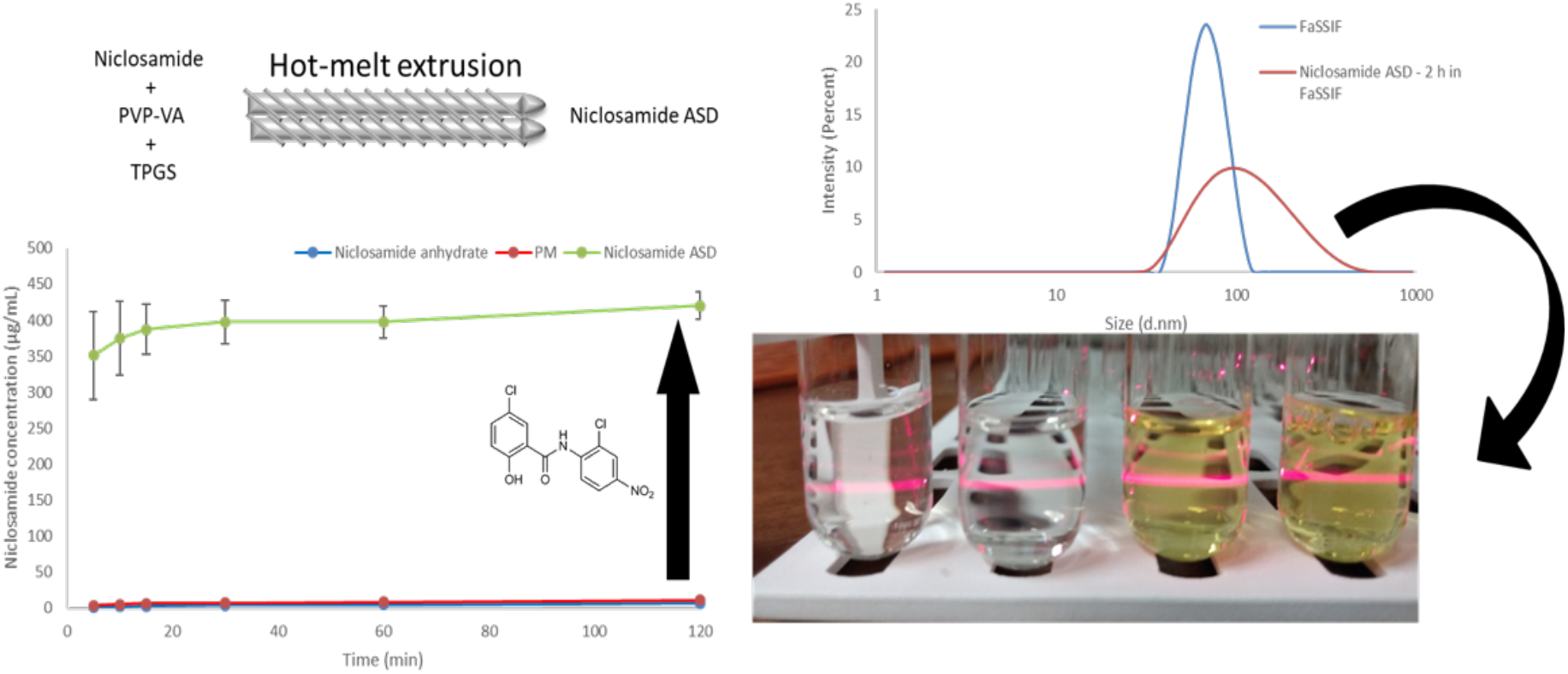

## 1. Introduction

*Niclosamide* is an FDA-approved anthelmintic drug that is one of the *Model List of Essential Medicines* produced by the World Health Organization (WHO) [1]. Some have advocated repurposing this drug for the treatment of various types of cancer and for use as a broad-spectrum antibacterial or antiviral drug, among other possible applications [2–5]. However, the repurposing of niclosamide is particularly challenging because it is a poorly water-soluble molecule and a poor glass former, which limits its oral absortion [6,7]. The water solubility of niclosamide has been reported to be about 13.32 μg/mL in its anhydrous form, but this falls to about 0.61 or 0.96 μg/mL for its monohydrate forms [8].

Several clinical trials have attempted to repurpose niclosamide. One of the clearest examples is the clinical trial NCT02532114, which was terminated early due to the low oral bioavailability of niclosamide, which exhibited Cmax values between 35.7 and 182 ng/mL after the oral administration of 500 mg three times daily. Furthermore, the authors stated that any attempt to repurpose the previously approved niclosamide product as a cancer therapy should be avoided, and efforts should be directed toward developing analogs that have higher bioavailability [9]. The efforts to increase niclosamide’s bioavailability include the use of co-crystals [7,10,11], solid lipid nanoparticles [12], dendrimer-like materials [13], micelles [14], nanosuspensions [15], nanoparticles [16], lipid emulsions [17], and nanocrystals [18].

The use of *amorphous solid dispersions* (ASDs) is a favorable formulation technique designed to increase the water solubility and bioavailability of poorly water-soluble drugs such as niclosamide. ASDs are, in general, dispersions in which the drug is dissolved in a solid matrix, usually a polymer [19]. These amorphous materials enhance apparent drug solubility by increasing the thermodynamic activity of the drug when it is molecularly and randomly dispersed in the polymer [20,21].

The tendency of a drug to become amorphous is evaluated using the concept of *glass forming ability* (GFA). The GFA of a drug falls into three categories: *class 1* (poor), *class 2* (modest), and *class 3* (good) [22]. Unfortunately, niclosamide is a GFA class 1 drug with a high propensity for recrystallization, which means it cannot form an amorphous solid on its own (i.e., a *neat* drug) [6]. However, niclosamide could form an ASD or a glass solution when it is dissolved in a solid matrix such as a polymer [23]. However, poor glass formers alone tend to recrystallize faster during dissolution [24].

In order to understand the mechanism of solubility and bioavailability improvement of niclosamide ASD, we evaluated the in vitro and in vivo performance of a niclosamide ASD prepared using hot-melt extrusion (HME). We assessed (1) the dissolution of the formulation in biorelevant media, (2) the polymer–niclosamide solid-state miscibility and intermolecular interactions, (3) the characterization of the nanoparticles generated by the ASD, (4) the in vitro permeation of the niclosamide ASD, and (5) the *in vivo* testing in a rat model using various strategies of administration.

## 2. Materials and Methods

### 2.1. Hot-melt extrusion (HME)

The polymer Kollindon® VA64 (PVP–VA) D-α-tocopheryl polyethylene glycol succinate (TPGS) was obtained from BASF, Germany. Niclosamide anhydrate was purchased from Shenzhen Nexconn Pharmatechs LTD (China). In the preparation of niclosamide ASD, a PVP–VA–niclosamide–TPGS blend in a 60:35:5 ratio was ground using a mortar and pestle until the mixture was homogeneous. Then, the mixture was processed using a HAAKE MiniLab II Micro Compounder (Thermo Electron Corporation, USA) set at 150 rpm and 180 °C. Thereafter, the extrudate was milled using a Tube Mill Control (IKA, Germany) and sieved to the range 45–125 μm.

### 2.2. HPLC analysis

The samples were measured at 331 nm using a Dionex HPLC system (Thermo Fisher Scientific Inc., USA) with a ZORBAX SB-C18 column (4.6 × 250 mm, 5 μm) (Agilent, USA) at a flow rate of 1 mL/min. Two mobile phases were used. The mobile phase A was a formic acid aqueous solution at 0.3%, and the mobile phase B was acetonitrile (Fisher Scientific, USA). They were mixed in a 40:60 ratio.

### 2.3. Dissolution testing

Dissolution tests of niclosamide ASD granules were conducted using FaSSIF (Biorelevant.com, United Kingdom) and the buffer (pH 6.5) required for its preparation. The FaSSIF medium was prepared according to the manufacturer’s specifications using the salts sodium hydroxide, monobasic sodium phosphate (Fisher scientific, USA), and sodium chloride (Sigma Aldrich, USA). The dissolution tests were performed in a Hanson SR8-Plus apparatus (Hanson Research Co., USA) using the 200 mL vessels and their paddles. Dissolution tests were conducted by adding the equivalent of 80 mg of niclosamide drug content into 150 mL of FaSSIF or buffer pH 6.5 media at 37.0 ± 0.5 °C and a paddle speed of 100 rpm. The sample points were measured using HPLC-UV at 5, 10, 15, 30, 60, and 120 min. When recollecting the samples, they were passed through polyethersulfone 0.2 μm filters. Then, 0.5 mL of the samples were mixed with 1 mL of acetone and 0.5 mL of acetonitrile for HPLC analysis.

The pH-shift dissolution tests were performed using the same equipment in two stages. First, 230 mg of niclosamide ASD was dissolved in 30 mL of HCl 0.01M for 30 min. Thereafter, the medium was changed to 150 mL of FaSSIF for 120 min.

### 2.4. Side-by-side diffusion cell

Side-by-side diffusion cells (PermeGear, USA) were employed to evaluate the diffusion of the niclosamide ASD through a 0.03 μm polyethersulfone membrane (Sterlitech Corp., USA). A similar method was used by Meng et al. (2019) [25]. The donor and receiver cells were filled with 34 mL of FaSSIF and decanol, respectively. We added 52.1 mg of the niclosamide ASD and 18.2 mg of niclosamide anhydrate to the donor cell at 37 °C and 850 rpm. The samples were collected from the receiver cell at 5, 10, 15, 30, 60, 120, and 180 min. Samples were measured using the same HPLC method described above.

### 2.5. Differential scanning calorimetry (DSC)

DSC was performed using a Model Q20 differential scanning calorimeter (TA Instruments, USA), increasing the temperature from 35 °C to 240 °C with a ramp temperature of 10 °C/min and a nitrogen purge of 50 mL/min.

### 2.6. Powder X-ray diffraction (XRD)

XRD studies were conducted using a Rigaku MiniFlex 600 II (Rigaku Americas, USA). The 2-theta angle was set at 5–40° (0.05° step, 2°/min, 40 kV, 15 mA).

### 2.7. Polarized light microscopy (PLM)

Samples were taken directly from the dissolution vessel and then placed on a glass slide. They were analyzed using a Olympus BX-53 polarized light microscope (Olympus Corporation of the Americas, USA) with a 200× objective and a QImaging QICAM digital camera (QImaging, Canada).

### 2.8. Particle size and zeta potential analysis

Samples were centrifuged at 13,000 rpm (14,300 rcf) for 10 min using a Microfuge®18 Centrifuge (Beckman Coulter, USA). The supernatant was measured using a Zetasizer Nano ZS (Malvern Instruments Ltd., UK).

### 2.9. Ultracentrifugation

To separate the particles and the unbound drug from the samples, an Airfuge™ Air-Driven Ultracentrifuge (Beckman Coulter, USA) was used at 30 psi for 30 min. Then, the supernatant was measured using HPLC.

### 2.10. Solid-state ^13^CNMR (ssNMR) spectroscopy

We performed ^13^CNMR spectroscopy using a Bruker AVANCE III HD 400 MHz spectrometer (Bruker, USA). One-dimensional ^13^C spectra were acquired using ramped cross-polarization of 70–100% on the ^1^H channel, with a magic angle spinning at 8 kHz and a total sideband suppression (TOSS) and high-power SPINAL64 proton decoupling. The acquisition parameters included a 2 ms contact time, a 60 s relaxation delay, and 256 scans.

### 2.11. Solution ^1^HNMR spectroscopy

^1^HNMR was conducted using an Agilent VNMRS 600 (600 MHz) (Agilent Technologies Inc, USA) at 25 °C. One dimensional ^1^H spectra were obtained using an acquisition time of 4 sec, 2-sec relaxation delay, and 64 scans. Samples were prepared by dissolving the drug, polymer, and drug–polymer mixtures in DMSO-d_6_. Initially, we tried to analyzed samples in D_2_0. However, the low solubility of niclosamide in aqueous environments generated poor quality NMRs. As Baghel *et al.* (2018), we selected DMSO because of its high dielectric constant, closer to water than other organic solvents that can solubilize niclosamide and PVP–VA. By doing this, we could get some idea of interactions between niclosamide and the polymers in a solution [26].

### 2.12. Animal studies

The oral pharmacokinetic analysis were conducted at Pharmaron (Ningbo, China). The study protocol was approved and conducted in accordance with the Institutional Animal Care and Use Committee (IACUC) guidelines at Pharmaron. (IACUC; Protocol Number AUP-2020-00060). In this study, niclosamide anhydrate and niclosamide ASD were administered to five rats per group (weight = 205.8 ± 2.9 g each) at a niclosamide dose of 10 mg/kg by oral gavage. The groups received a FaSSIF suspension of niclosamide anhydrate at 1.5 mg/mL, a FaSSIF suspension of niclosamide ASD at 1.5 mg/mL, and size 9 mini capsules (Braintree Scientific, USA) containing niclosamide ASD, respectively (Three groups in total). In this last group, the capsule size 9 contained 60% niclosamide ASD, 15% EXPLOTAB^©^, and 25% sodium bicarbonate.

The samples were measured using an AB Sciex Triple Quad 5500 LC/MS/MS with an Agilent Eclipse XDB-C18 column (2.1 × 150 mm, 5 μm) (Agilent, USA) at a flow rate of 0.6 mL/min. Two mobile phases were used. The mobile phase A was a 0.1% formic acid aqueous solution, and the mobile phase B was a mixture of 5% water and 95% acetonitrile (0.1% formic acid). They were mixed as shown in the supplemental materials (Table S1). Then, 50 μL of plasma with 5 μL of methanol were added to 200 μL of methanol containing an internal standard mixture for protein precipitation. The samples were vortexed for 30 s and underwent centrifugation for 15 min at 4,000 rpm and 4 °C. Thereafter, the supernatant was diluted three times with water, and 2 μL were injected into the HPLC.

### 2.13. Statistical analyses

Student t-test (*p* < .05) was used for statistical analysis when comparing two groups. One-way ANOVA analysis (*p* < .05) with post hoc Tukey’s multiple comparison test was used when comparing three groups. The analyses were performed using JMP Pro 15 software.

## 3. Results

### 3.1. Hot-melt extrusion successfully prepared amorphous extrudates of niclosamide

Niclosamide ASD formulations were characterized using DSC and XRD to confirm their amorphous nature. Figure 1A depicts niclosamide’s solid dispersion as amorphous, based on the lack of diffraction peaks associated with the crystalline niclosamide anhydrate material. Moreover, the thermogram of niclosamide ASD did not exhibit the melting point of the drug at 230 °C (Figure 1A). However, the niclosamide’s melting endotherm was not observed in physical mixture blends of niclosamide and PVP–VA, as a result of the drug dissolving in the polymer before reaching 230 °C (data not shown). As such, the niclosamide was considered amorphous at the limit of XRD sensitivity. Thus, niclosamide dissolved at a molecular level in the PVP–VA matrix during HME manufacturing.

**Figure 1.**
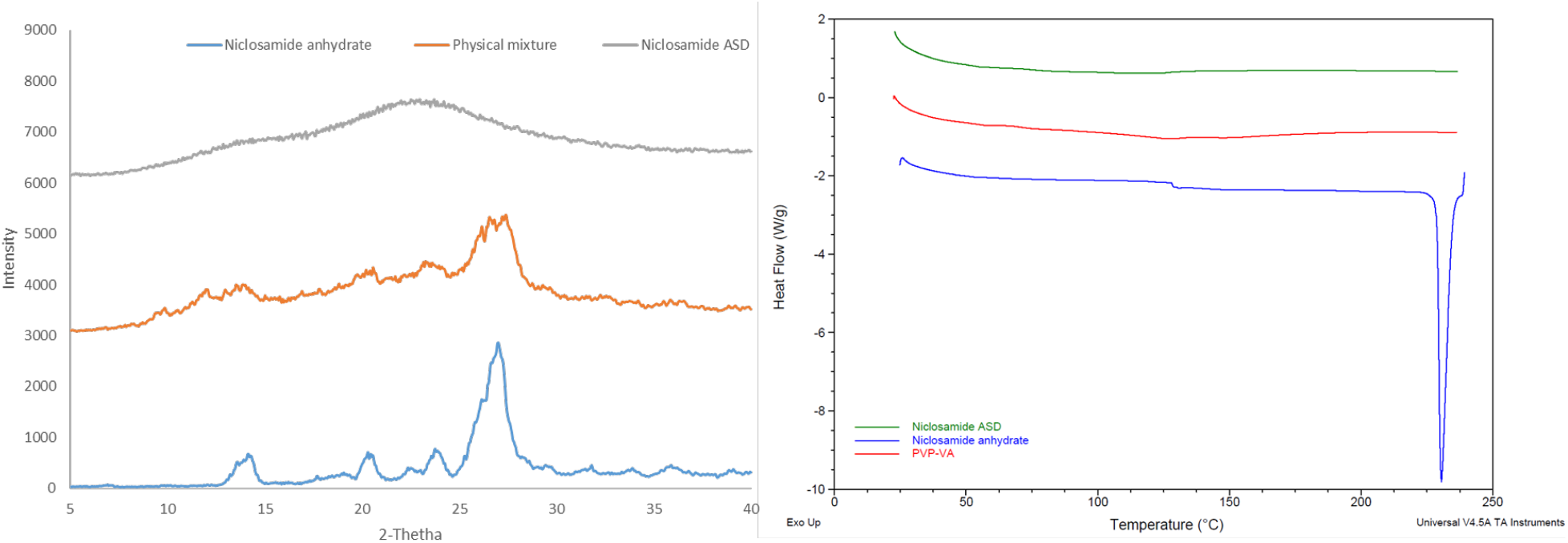
(A) XRD diffraction pattern. (B) DSC thermogram of niclosamide ASD (extrudate), its physical mixture, and niclosamide anhydrate

### 3.2. The amorphous extrudates increased niclosamide’s apparent solubility

The dissolution tests using the biorelevant FaSSIF media showed a notable increase in the apparent solubility of niclosamide, as shown in Figure 2A. Notably, the granules of the niclosamide ASD reached a plateau more than 60 times higher than the crystalline niclosamide anhydrate after 2 h of dissolution testing (420.2 ± 19.0 vs. 6.6 ± 0.4 μg/mL, *p* < .001). This plateau was found to persist for more than 24 h, even with some signs of crystallization under PLM (Figure S1). As the formulation design screenings were performed using the solvent shift methods and 0.2 μm filters, it was relevant to measure the samples using DLS to identify the existence of nanoparticles in the system. The Zetasizer Nano ZS was able to detect nanoparticles with a mean particle size of about 100 nm and a zeta potential of −13.6 ± 1.0 mV, probably due to the FaSSIF (Figure 2B). Evidently, this formulation was able to enhance the apparent solubility 60-fold by generating nanoparticles that could not be removed by the filter. After these findings, ultracentrifugation was used to concentrate the particles and measure the supernatant. Surprisingly, the niclosamide concentration was just 11.4 ± 8.5 μg/mL after being subjected to the dissolution test for 120 min, which is lower than the 2-fold increase when compared with the anhydrate in FaSSIF (measured after filtration).

**Figure 2.**
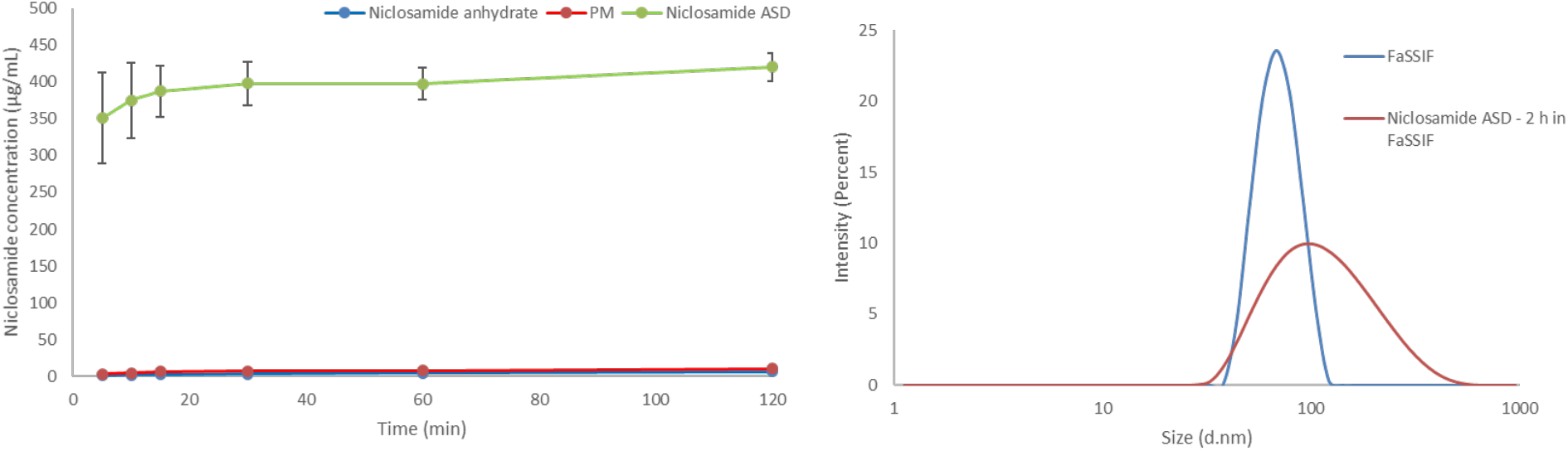
The dissolution profile of niclosamide ASD, its physical mixture (PM), and niclosamide anhydrate in FaSSIF medium. The samples were taken and passed through 0.2 μm filters.

In summary, the apparent increase in niclosamide solubility from the ASD was largely attributable to the generation of nanoparticles as opposed to increased drug supersaturation. Side-by-side diffusion cells were used to evaluate both dissolution and permeation of the drug across a membrane in order to determine whether the apparent solubility increase resulted from the presence of the nanoparticles can increase niclosamide’s diffusion.

### 3.3. The amorphous extrudates increased niclosamide diffusion in side-by-side diffusion cells

Side-by-side diffusion cells confirmed that the increase in apparent solubility translates to increased diffusion of niclosamide through a membrane into the receiver cell (decanol), as shown in Figure 3. The niclosamide ASD formulation achieved a concentration of 53.8 ± 13.1 μg/mL. In contrast, niclosamide anhydrate reached a concentration of only 12.1 ± 1.3 μg/mL in the receiver cell (p = .005).

**Figure 3.**
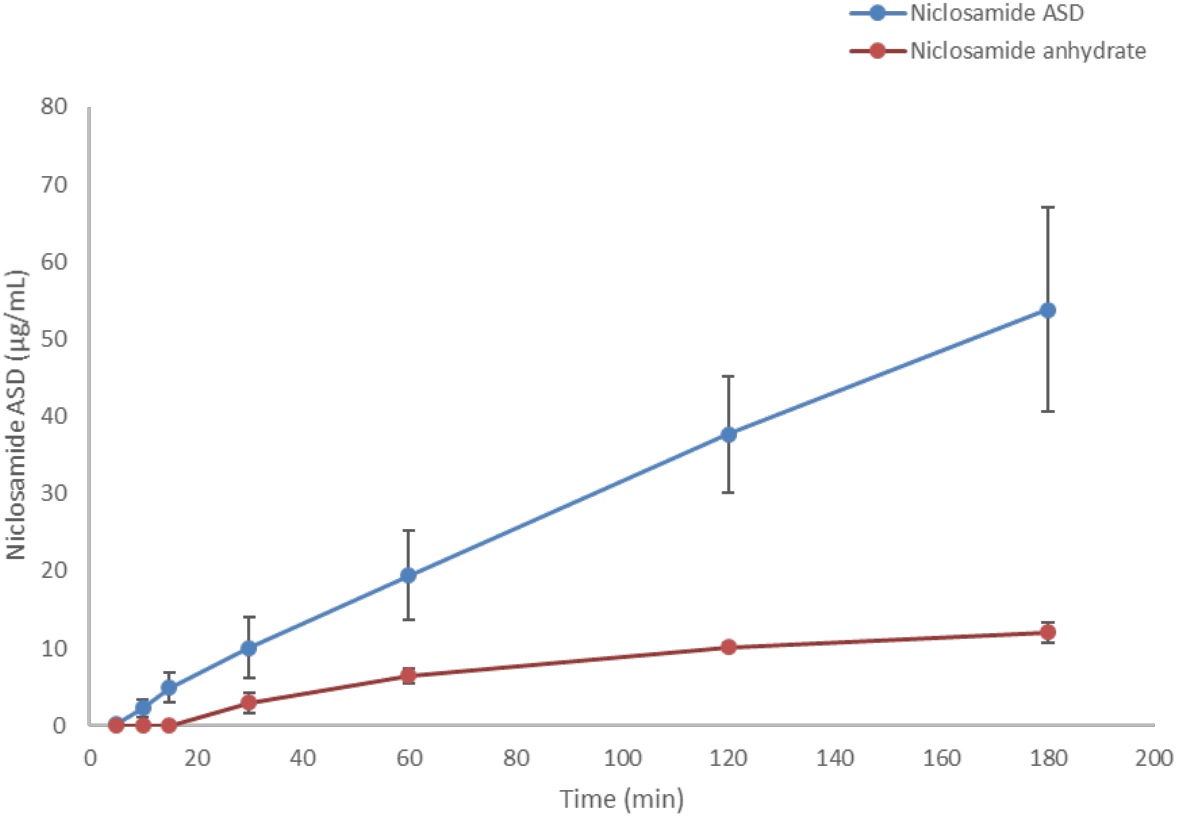
Diffusion profiles of niclosamide ASD and niclosamide anhydrate. The donor and receiver cells were filled with FaSSIF and decanol, respectively.

### 3.4. FTIR

The peaks of niclosamide and PVP–VA were assigned as described in previous studies [7,27–29]. As can be seen in Figure 4, noticeable changes were observed when comparing niclosamide ASD and its physical mixture throughout the spectra. These differences indicate interactions between niclosamide and PVP–VA [30]. One example of these interactions can be seen in the C=O stretchings of niclosamide and PVP–VA, which are highlighted in the figure with dotted lines. It is known that the 2-pyrrolidinone group acts as a hydrogen bond acceptor group capable of stabilizing -NH and -OH hydrogen bond donors, moieties present in niclosamide [31]. Interestingly, this PVP–VA carbonyl stretching (amide) even shifted in the physical mixture. This indicates a strong interaction, a phenomenon that has been seen in other studies with niclosamide and other excipients [27]. Solidstate NMR was conducted to gather further evidence about these interactions between niclosamide and PVP–VA.

**Figure 4.**
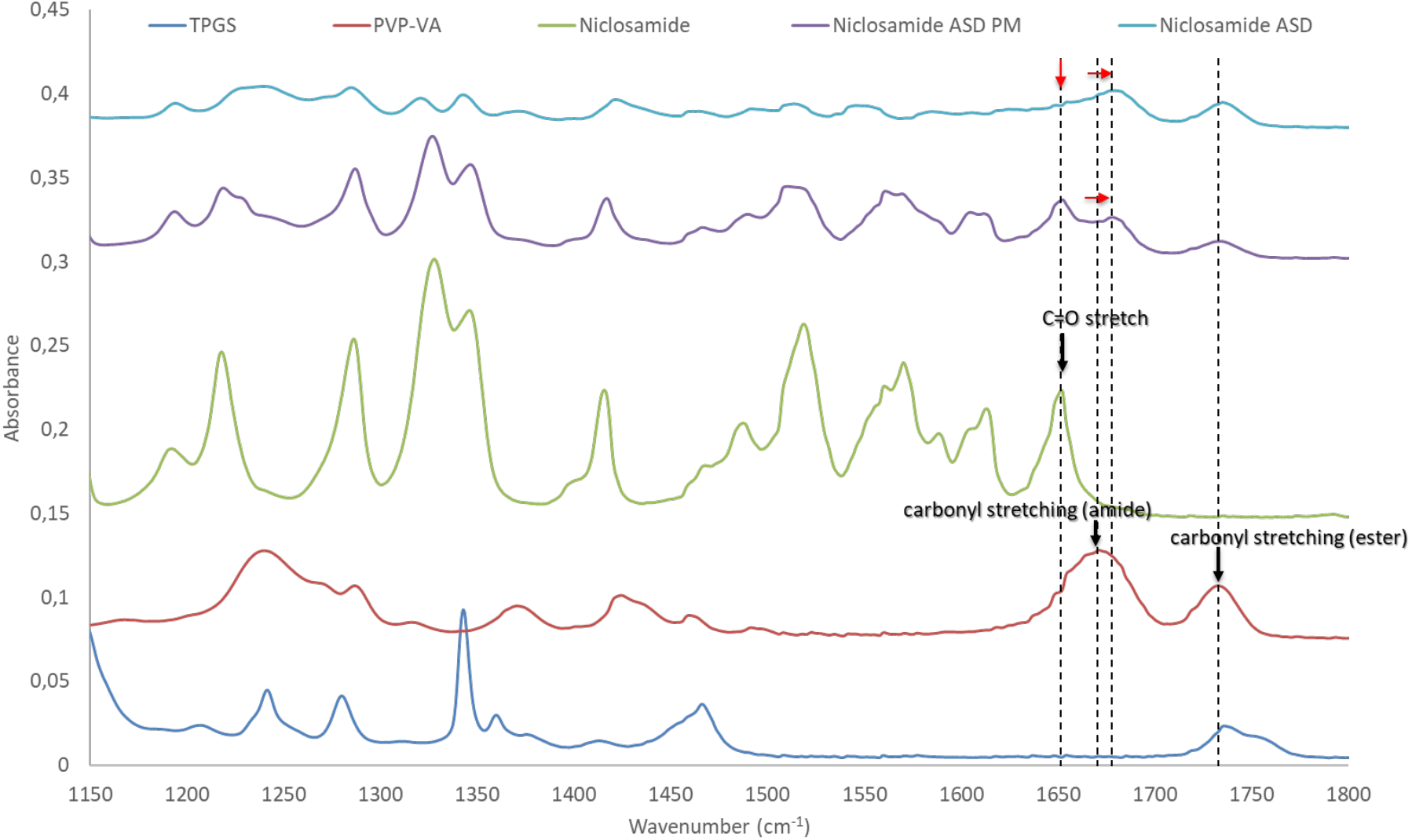
FTIR spectra of niclosamide ASD, its physical mixture, and its constituents. The carbonyl stretchings are highlighted with dotted lines. Red arrows indicate their peak shifts.

### 3.5. Solid-state NMR shows the importance of the phenolic group in the amorphous dispersion

We used ssNMR to analyze niclosamide anhydrate, PVP–VA, and an amorphous extrudate at 35% DL (without TPGS) as well as its physical mixture. The carbon peaks of the ssNMR were assigned according to previous literature [10,11]. Noticeably, the niclosamide phenolic carbon showed a downfield shift in the physical mixture and even more so in the extrudate (i.e., the glass material) (Figure **Error! Reference source not found.**5). Moreover, a broadening peak was observed in the niclosamide ASD sample. Peak broadening is characteristic of reduced or eliminated crystalline structure due to a higher disorder of the drug–polymer interactions [32]. Figure 5 shows that niclosamide aromatic hydrogens were disrupted by the presence of the polymer due to peak broadening, which confirms observations from FTIR (i.e., the ASD versus the physical mixture) [32].

**Figure 5.**
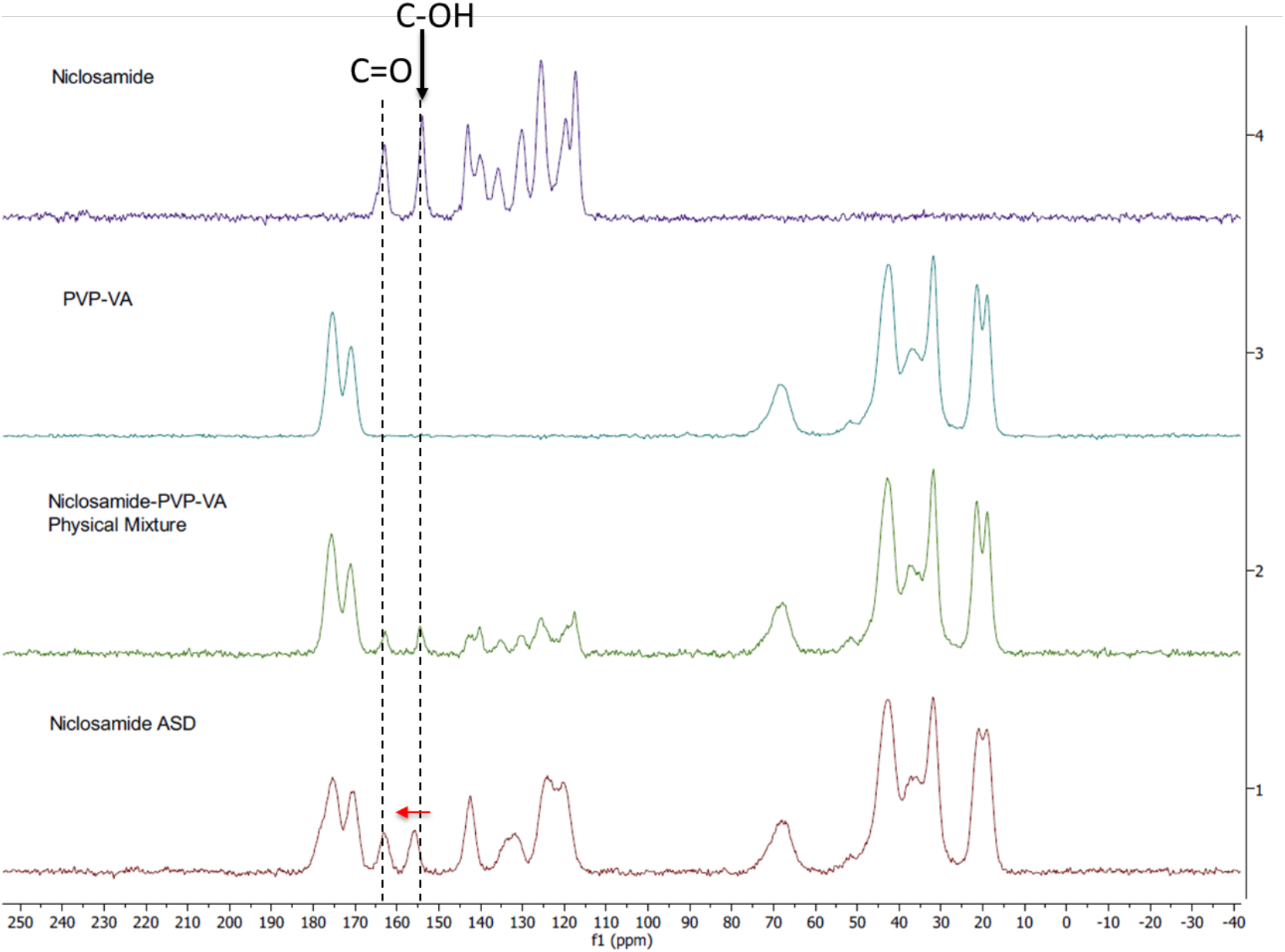
Solid-state ^13^CNMR spectra of niclosamide anhydrate, PVP–VA, 35% niclosamide-65% PVP–VA physical mixture and extrudate. The phenolic and carbonyl carbons are highlighted with dotted lines. The red arrow indicates the peak shift of phenolic carbon.

Interestingly, the chemical shift (dotted line) involving the phenolic carbon could be pH dependent in a water solution. Experimental confirmation is needed.

### 3.6. Solution NMR showed the relevance of the 2-pyrrolidinone group for niclosamide’s stabilization after dissolution

Due to niclosamide’s poor water solubility, DMSO-d_6_ was used for qualitative purposes instead of D_2_O. Initially, we screened several polymers, such as PVP and PVP–VA, for the development of the niclosamide ASD formulation (without TPGS). PVP–VA was observed to be superior in increasing the apparent solubility of niclosamide (data not shown). We hypothesized that the vinyl acetate groups (VA) in PVP–VA were mainly responsible for these differences, as was observed in another drug–polymer system previously reported [33].

To test our hypothesis, we conducted ^1^HNMR for both systems (i.e., the niclosamide–PVP and the niclosamide–PVP–VA) to see whether the VA groups play a key role in interacting with the drug. The solution ^1^HNMR showed interactions between niclosamide and the polymers (Figure 6). Interestingly, the more noticeable downfield shifts were seen in the niclosamide–PVP sample. According to Ueda *et al.* (2020), peak broadening is related to mobility suppression [34]. This means that, contrary to our previous dissolution observations, PVP interacts more strongly with niclosamide than with PVP–VA in solution. Once again, there is a shift of the phenolic -OH (smaller than -NH group); this interaction probably plays a significant role in aqueous environments due to ionization (pKa = 6.89). In the case of the niclosamide–PVP–VA, there was a small upfield shift in the amide hydrogen from niclosamide. Kawakami *et al.* (2018) observed that even the most favorable interaction could be secondary to avoid the drug’s crystallization if the polymer–drug system does not disintegrate or fails to dissolve properly [35]. This seems to be true in the case of the PVP–niclosamide ASD, because we observed differences in wettability while conducting the dissolution tests using niclosamide–PVP and niclosamide–PVP–VA extrudates (data not shown).

**Figure 6.**
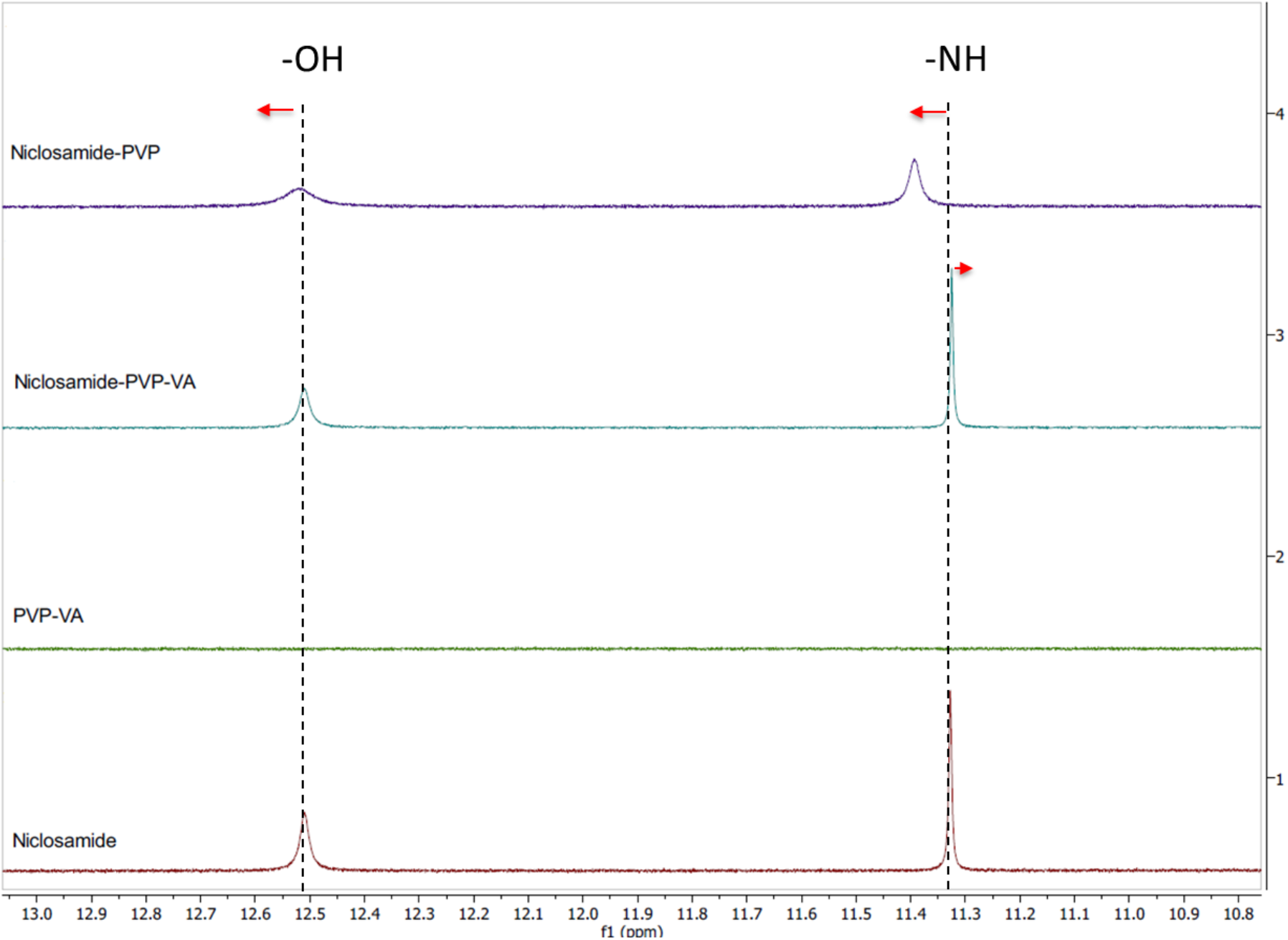
Solution ^1^HNMR spectra of niclosamide anhydrate, PVP, PVP–VA, and a mixture of 35% niclosamide and 65% PVP–VA. The phenolic and amide hydrogens are highlighted with dotted lines. The red arrows indicate peak shifts of phenolic and amide hydrogen.

### 3.7. pH shift dissolution testing confirms that the amorphous extrudates crystallize in acidic conditions

NMR and FTIR showed the relevance of niclosamide’s phenolic group in the ASD, which may ionize at gastrointestinal pH levels. A dissolution test with pH shifting was performed to determine the behavior of the niclosamide ASD at pH 2 for 30 min, then at pH 6.5 (in FaSSIF medium) for 120 min. The dissolution results show a major reduction in apparent solubility, even after the pH shift with FaSSIF. PLM confirms that the niclosamide ASD crystallized under acidic conditions (Figure 7). Even after the pH shift, the niclosamide ASD could not achieve the increase in apparent solubility that was seen in the FaSSIF medium alone (Figure S2). This reinforces the concerns raised by spectroscopy and confirms the importance of the phenolic group interaction with the polymer. We propose that this is due to niclosamide’s pKa and its changes in solubility. Our results support the limitations of conducting a pharmacokinetic study in animals due to the lack of enteric-coated options to administer the formulations.

**Figure 7.**
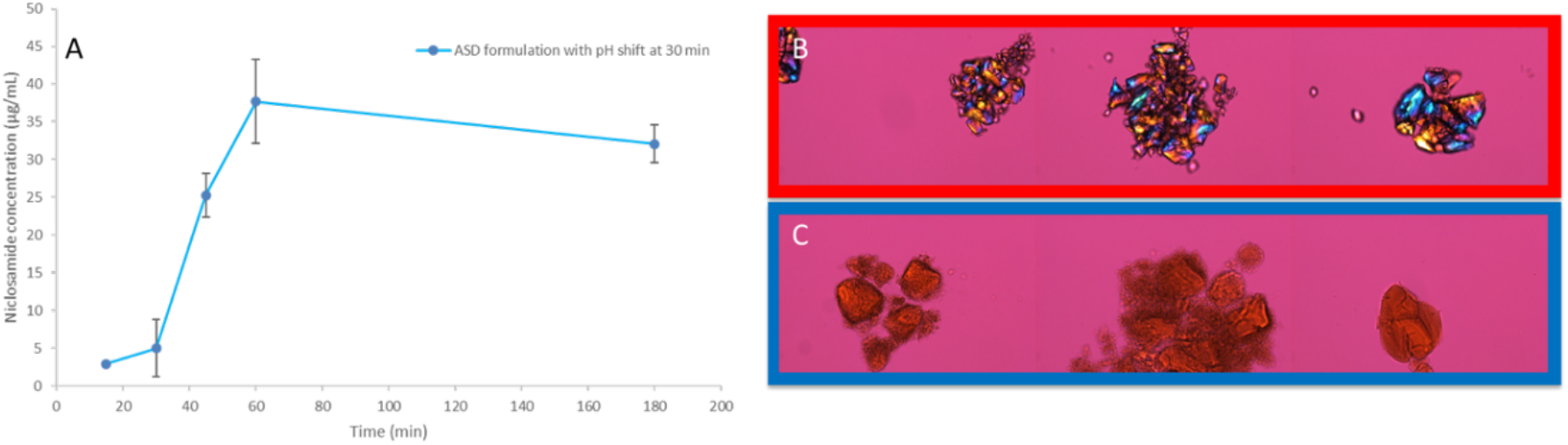
(A) pH-shift dissolution test of niclosamide ASD. (B) PLM images at 200× (30 min). The dotted line shows the pH-shift at 30 min. (C) PLM of niclosamide ASD at 120 min without pH shift. Signs of crystallization were not observed.

### 3.8. The amorphous extrudates increased the bioavailability of niclosamide

The method of administration of niclosamide was designed in such a way that it considered the limitation of using a rat model. McConnell *et al.* (2008) previously reported an in-depth study of the gastrointestinal tract of rats, and they found that the rat stomach has a pH of 3.2 ± 1.0 and 3.9 ± 1.0 in the fed and fasted state, respectively [36]. The pH level of the rat intestine never exceeds 6.6. They concluded that rats are not a suitable model for the study of pH-sensitive dosage forms that require pH values that mimic the human gastrointestinal tract [36]. We worked to overcome these limitations by administering an oral gavage of the formulation already dispersed in FaSSIF with double the buffer strength in order to neutralize the rat stomach pH. In the experiment, the salts and FaSSIF comprised the suspension medium used to disperse the niclosamide ASD by vortexing.

The niclosamide ASD administered using the 2× buffer capacity FaSSIF suspension and the capsules (containing sodium bicarbonate and EXPLOTAB®) achieved similar bioavailability (see Table 1 and Figure 8, *p* > .75). We observed a more than 2-fold increase in bioavailability from the niclosamide ASD suspension compared to niclosamide anhydrate (*p* < .05). As expected, the capsules increased the Tmax because the formulation underwent disintegration and dissolution. The administration of the niclosamide ASD using capsules did not achieve a statistically significant increase in bioavailability (*p* = .15).

**Table 1.** Pharmacokinetic parameter profiles (in rats) of niclosamide anhydrate suspended in FaSSIF, niclosamide ASD suspended in FaSSIF, and niclosamide ASD in capsules (n = 5).

| PK parameters | Niclosamide anhydrate suspension in FaSSIF | Niclosamide ASD suspension in FaSSIF | Niclosamide ASD in capsules |
| --- | --- | --- | --- |
| $T_{1/2}$ (h) | 1.00 (0.30) | 1.59 (1.34) | 0.84 (0.01) |
| $T_{max}$ (h) | 3.60 (0.89) | 2.40 (1.52) | 4.40 (0.89) |
| $C_{max}$ (ng/mL) | 48.3 (20.6) | 123 (56) | 122 (71) |
| $AUC_{last}$ (h*ng/mL) | 168 (64) | 398 (115) | 338 (193) |
| $AUC_{Inf}$ (h*ng/mL) | 188 (84) | 495 (239) | 463 (224) |
| $AUC_{\%Extrap\_obs}$ (%) | 8.4 (7.0) | 12.2 (21.4) | 7.98 (0.48) |
| $MRT_{Inf\_obs}$ (h) | 3.56 (0.70) | 3.71 (2.20) | 4.04 (0.12) |
| $AUC_{last/D}$ (h*mg/mL) | 16.8 (6.4) | 39.8 (11.5) | 33.8 (19.3) |

**Figure 8.**
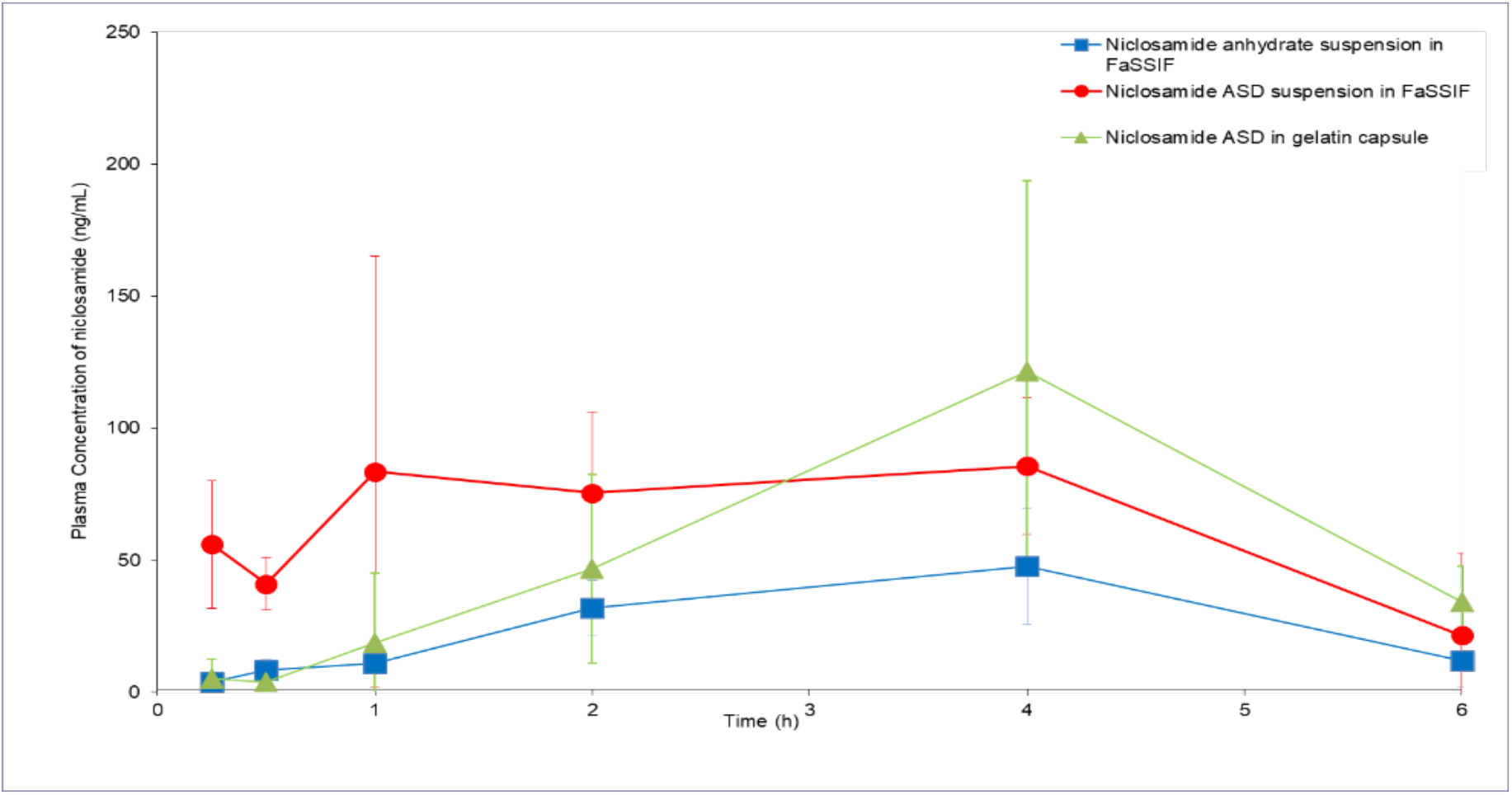
Pharmacokinetic profiles (in rats) of niclosamide anhydrate suspended in FaSSIF, niclosamide ASD suspended in FaSSIF, and niclosamide ASD in capsules (n = 5).

## 4. Discussion

The selection of the formulation composition was made using the solvent-shift method and FaSSIF, as described by Palmelund et al. (2017), but employing 0.2 μm filters for quantification in order to measure an apparent solubility that included the presence of nanoparticles [37]. It is common to find studies in the literature that report dissolution testing using HPLC- or UV-Vis-based methodologies after a filtration step, similar to our present study [38,39].

It is known that some ASDs can generate nanometric drug-rich colloids, which act as drug reservoirs for fast dissolution [20,35]. These colloids can lead to a confounding supersaturation measurement above the amorphous drug’s solubility if they are small enough to pass through filters and resist precipitation after conventional centrifugation [40]. In our development of a niclosamide ASD, we acknowledged the potential confounding factor of nanoparticles, and we used this to create a formulation that could avoid the crystallization of niclosamide, which is a poor glass former.

Moreover, there is evidence that ASDs that generate drug-rich colloids can increase bioavailability compared to ASDs that do not form them [41]. The generation of nanoparticles can be misleading when determining the supersaturation advantage. However, they can function as drug reservoirs that maintain a supersaturated concentration that drives absorption. These nanoparticles can also reduce the tendency to crystallize by inhibiting drug nucleation due to specific polymer–drug intermolecular interactions [42].

The formation of nanoparticles depends on the presence of FaSSIF as the bile salts serving as a stabilizer reduce the mean particle size and confer an adequate zeta potential for electrostatic stabilization [43,44]. To our surprise, this suspension remained stable for long periods of time. This drug behavior is similar to what was previously described as the “spring and hover” effect, in which a noticeable increase in apparent solubility reaches a plateau and persists over a long period of time [39]. Interestingly, in their work with nicotinamide–ibuprofen co-crystals, they achieved a 70-fold increase in supersaturation, and they measured the drug supersaturation using 0.22 μm filters (similar to our work).

Several studies show that specific drug–polymer interactions are critical for increasing supersaturation, apparent drug solubility, and even solid-state miscibility/stability. FTIR and ssNMR showed changes in all niclosamide signals, including the physical mixture and the extrudates (i.e., the glassy material). This confirms intimate interactions between niclosamide and the polymer. The ssNMR in particular showed not only peak broadening but also a phenolic carbon shift, which indicates specific molecular interactions after ionization at intestinal pH (pKa of niclosamide = 6.89).

It was not possible to obtain a suitable ^1^HNMR using D_2_O as a solvent with all the components of the composition (i.e., the niclosamide ASD and FaSSIF). Therefore, we selected DMSO as a solvent for qualitative purposes because its dielectric constant is closer to water than other organic solvents. Interestingly, this experiment showed that, in solution, the 2-pyrrolidinone group from PVP–VA seems to be critical for niclosamide stabilization. These groups are good hydrogen acceptors, especially for the -OH and the -NH groups of niclosamide that experienced chemical shifting and peak broadening.

These results suggest a potential threat for the oral administration of the formulation due to pH changes in the stomach. Niclosamide is a BCSII and weakly acidic drug that remains mostly un-ionized at stomach pH levels. The dissolution data with a pH shift confirm that our concerns about crystallization were warranted, as shown in Figure 7. Kawakami et al (2018). encountered a similar situation when working with fenofibrate ASDs using various polymers in acidic media. They explained that these polymers dissolved and left behind drug-rich domains that crystallized [35].

In our pharmacokinetic study in rats, we acknowledged the challenge of the gastrointestinal pH in rats. We administered the formulations as a suspension using FaSSIF and as capsules containing sodium bicarbonate to counteract the acid pH of the stomach. The niclosamide ASD suspension resulted in an AUC_last_ (last time point) more than double the administration of niclosamide anhydrate suspension (Table 1). Administration by capsule did not achieve a statistically significant increase in bioavailability.

It is important to note that all our initial studies were conducted with the intent of simulating the human fasting intestine environment using biorelevant media in terms of pH, osmolarity, and bile salts [45]. Unfortunately, rats have different gastrointestinal pH values than humans, and reliable enteric-coated capsules are not commercially available for administering pH-sensitive formulations. The differences in pH and bile salt concentrations between the rat model and in vitro testing can generate substantial discrepancies in formulation performance, rendering it difficult to predict [46].

When contrasted with the literature, the performance of the niclosamide ASD was similar to the administration of dissolved niclosamide using DMSO–cremophor EL–water mixtures (429 ± 100 ng/mL · h) [47]. Dosing solubilized niclosamide (a BCS class II molecule with a stable crystalline structure) using solvents or oils increased bioavailability because the drug was already available for oral absortion in molecular form [1,48,49]. Based on our results, a niclosamide ASD should now be formulated as an enteric-coated dosage form to protect it from gastric acid and subsequent crystallization.

## 5. Conclusions

This study demonstrates that an amorphous solid dispersion of niclosamide increased the drug’s bioavailability in a rat model. This model was particularly challenging for the formulation due to the differences between the in vitro and in vivo models. The results indicate that the repurposing of niclosamide as an oral dosage form is viable, and a greater increase in bioavailability is expected if the drug is formulated as an enteric-coated product. Overall, it is feasible to use HME manufacturing to increase niclosamide’s bioavailability. This will pave the way for new applications of the drug as an antibacterial/antiviral or as an oral therapy for cancer or COVID-19, among others.

## Acknowledgments

We thank Dr. Steve Sorey and Dr. Garrett Blake for their help with the NMR spectra.

## Supplementary materials

**Table S1.** Mobile phase gradient that was used for analyzing plasma sample.

| Time (min) | A (%) | B (%) |
| --- | --- | --- |
| 0.20 | 85.0 | 15.0 |
| 2.00 | 50.0 | 50.0 |
| 2.50 | 50.0 | 50.0 |
| 4.00 | 0.00 | 100 |
| 4.50 | 0.00 | 100 |
| 4.51 | 85.0 | 15.0 |
| 5.00 | 85.0 | 15.0 |

**Figure S1.**
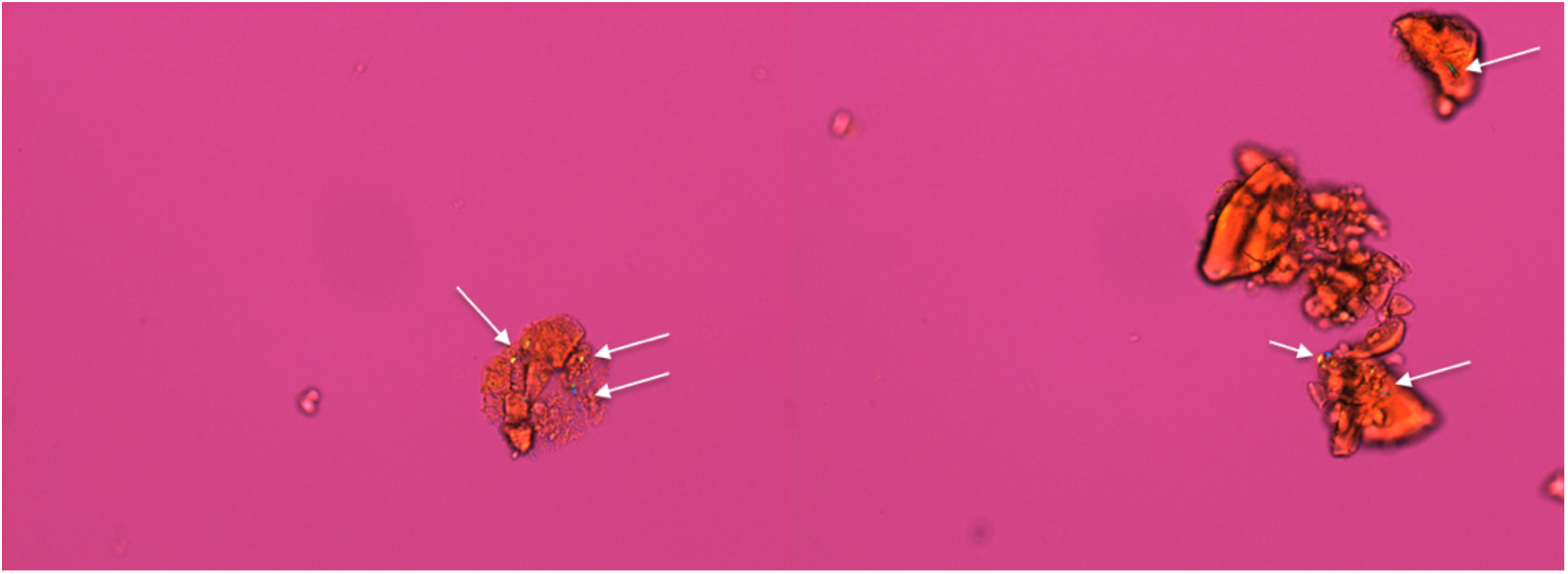
PLM of niclosamide ASD at 24 h without pH-shift. Signs of crystallization were observed (white arrows).

**Figure S2.**
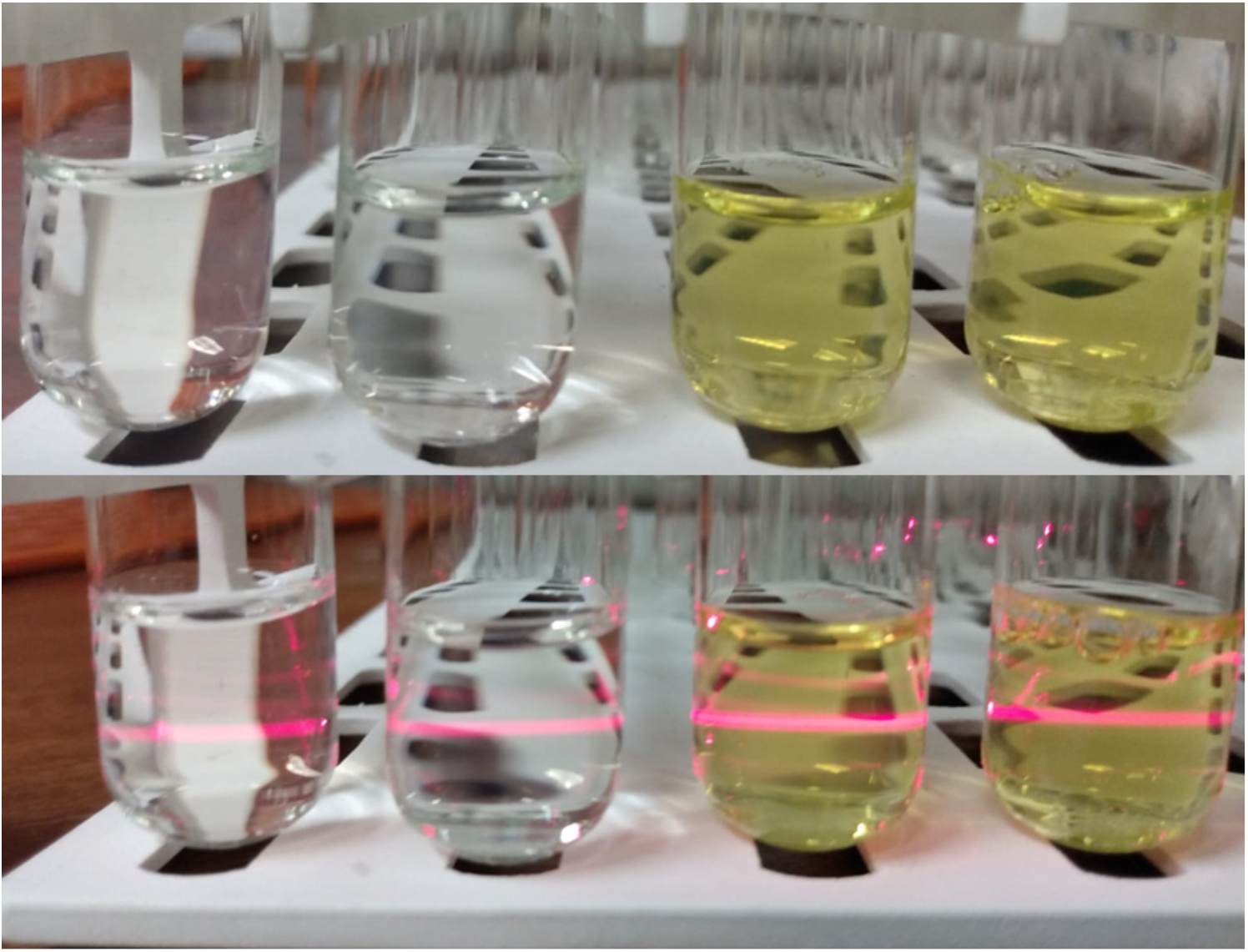
The image shows the appearance of samples for HPLC after filtration. The samples in the right are with pH-shift (transparent) and in the left without pH-shift after 24 h. The laser beam shows the presence of colloidal species.

## References

1. Barbosa, E.J.; Löbenberg, R.; de Araujo, G.L.B.; Bou-Chacra, N.A. Niclosamide Repositioning for Treating Cancer: Challenges and Nano-Based Drug Delivery Opportunities. Eur. J. Pharm. Biopharm. Off. J. Arbeitsgemeinschaft Pharm. Verfahrenstechnik EV 2019, 141, 58–69, doi:10.1016/j.ejpb.2019.05.004.

2. Xu, J.; Shi, P.-Y.; Li, H.; Zhou, J. Broad Spectrum Antiviral Agent Niclosamide and Its Therapeutic Potential. ACS Infect. Dis. 2020, doi:10.1021/acsinfecdis.0c00052.

3. Li, Y.; Li, P.-K.; Roberts, M.J.; Arend, R.C.; Samant, R.S.; Buchsbaum, D.J. Multi-Targeted Therapy of Cancer by Niclosamide: A New Application for an Old Drug. Cancer Lett. 2014, 349, 8–14, doi:10.1016/j.canlet.2014.04.003.

4. Chen, W.; Mook, R.A.; Premont, R.T.; Wang, J. Niclosamide: Beyond an Antihelminthic Drug. Cell. Signal. 2018, 41, 89–96, doi:10.1016/j.cellsig.2017.04.001.

5. Tam, J.; Hamza, T.; Ma, B.; Chen, K.; Beilhartz, G.L.; Ravel, J.; Feng, H.; Melnyk, R.A. Host-Targeted Niclosamide Inhibits C. Difficile Virulence and Prevents Disease in Mice without Disrupting the Gut Microbiota. Nat. Commun. 2018, 9, 1–11, doi:10.1038/s41467-018-07705-w.

6. Alhalaweh, A.; Alzghoul, A.; Kaialy, W.; Mahlin, D.; Bergström, C.A.S. Computational Predictions of Glass-Forming Ability and Crystallization Tendency of Drug Molecules. Mol. Pharm. 2014, 11, 3123–3132, doi:10.1021/mp500303a.

7. Sanphui, P.; Kumar, S.S.; Nangia, A. Pharmaceutical Cocrystals of Niclosamide. Cryst. Growth Des. 2012, 12, 4588–4599, doi:10.1021/cg300784v.

8. van Tonder, E.C.; Maleka, T.S.P.; Liebenberg, W.; Song, M.; Wurster, D.E.; de Villiers, M.M. Preparation and Physicochemical Properties of Niclosamide Anhydrate and Two Monohydrates. Int. J. Pharm. 2004, 269, 417–432, doi:10.1016/j.ijpharm.2003.09.035.

9. Schweizer, M.T.; Haugk, K.; McKiernan, J.S.; Gulati, R.; Cheng, H.H.; Maes, J.L.; Dumpit, R.F.; Nelson, P.S.; Montgomery, B.; McCune, J.S.; et al. A Phase I Study of Niclosamide in Combination with Enzalutamide in Men with Castration-Resistant Prostate Cancer. PLoS ONE 2018, 13, doi:10.1371/journal.pone.0198389.

10. Grifasi, F.; Chierotti, M.R.; Gaglioti, K.; Gobetto, R.; Maini, L.; Braga, D.; Dichiarante, E.; Curzi, M. Using Salt Cocrystals to Improve the Solubility of Niclosamide. Cryst. Growth Des. 2015, 15, 1939–1948, doi:10.1021/acs.cgd.5b00106.

11. Luedeker, D.; Gossmann, R.; Langer, K.; Brunklaus, G. Crystal Engineering of Pharmaceutical Co-Crystals: “NMR Crystallography” of Niclosamide Co-Crystals. Cryst. Growth Des. 2016, 16, 3087–3100, doi:10.1021/acs.cgd.5b01619.

12. Rehman, M.U.; Khan, M.A.; Khan, W.S.; Shafique, M.; Khan, M. Fabrication of Niclosamide Loaded Solid Lipid Nanoparticles: In Vitro Characterization and Comparative in Vivo Evaluation. Artif. Cells Nanomedicine Biotechnol. 2018, 46, 1926–1934, doi:10.1080/21691401.2017.1396996.

13. Xie, Y.; Yao, Y. Octenylsuccinate Hydroxypropyl Phytoglycogen Enhances the Solubility and In-Vitro Antitumor Efficacy of Niclosamide. Int. J. Pharm. 2018, 535, 157–163, doi:10.1016/j.ijpharm.2017.11.004.

14. Russo, A.; Pellosi, D.S.; Pagliara, V.; Milone, M.R.; Pucci, B.; Caetano, W.; Hioka, N.; Budillon, A.; Ungaro, F.; Russo, G.; et al. Biotin-Targeted Pluronic® P123/F127 Mixed Micelles Delivering Niclosamide: A Repositioning Strategy to Treat Drug-Resistant Lung Cancer Cells. Int. J. Pharm. 2016, 511, 127–139, doi:10.1016/j.ijpharm.2016.06.118.

15. Costabile, G.; d’Angelo, I.; Rampioni, G.; Bondì, R.; Pompili, B.; Ascenzioni, F.; Mitidieri, E.; d’Emmanuele di Villa Bianca, R.; Sorrentino, R.; Miro, A.; et al. Toward Repositioning Niclosamide for Antivirulence Therapy of Pseudomonas Aeruginosa Lung Infections: Development of Inhalable Formulations through Nanosuspension Technology. Mol. Pharm. 2015, 12, 2604–2617, doi:10.1021/acs.molpharmaceut.5b00098.

16. Naqvi, S.; Mohiyuddin, S.; Gopinath, P. Niclosamide Loaded Biodegradable Chitosan Nanocargoes: An in Vitro Study for Potential Application in Cancer Therapy. R. Soc. Open Sci. 4, 170611, doi:10.1098/rsos.170611.

17. Zhang, X.; Zhang, Y.; Zhang, T.; Zhang, J.; Wu, B. Significantly Enhanced Bioavailability of Niclosamide through Submicron Lipid Emulsions with or without PEG-Lipid: A Comparative Study. J. Microencapsul. 2015, 32, 496–502, doi:10.3109/02652048.2015.1057251.

18. Ye, Y.; Zhang, X.; Zhang, T.; Wang, H.; Wu, B. Design and Evaluation of Injectable Niclosamide Nanocrystals Prepared by Wet Media Milling Technique. Drug Dev. Ind. Pharm. 2015, 41, 1416–1424, doi:10.3109/03639045.2014.954585.

19. Schittny, A.; Huwyler, J.; Puchkov, M. Mechanisms of Increased Bioavailability through Amorphous Solid Dispersions: A Review. Drug Deliv. 2019, 27, 110–127, doi:10.1080/10717544.2019.1704940.

20. Indulkar, A.S.; Lou, X.; Zhang, G.G.Z.; Taylor, L.S. Insights into the Dissolution Mechanism of Ritonavir–Copovidone Amorphous Solid Dispersions: Importance of Congruent Release for Enhanced Performance. Mol. Pharm. 2019, 16, 1327–1339, doi:10.1021/acs.molpharmaceut.8b01261.

21. Thakkar, R.; Thakkar, R.; Pillai, A.; Ashour, E.A.; Repka, M.A. Systematic Screening of Pharmaceutical Polymers for Hot Melt Extrusion Processing: A Comprehensive Review. Int. J. Pharm. 2020, 576, 118989, doi:10.1016/j.ijpharm.2019.118989.

22. Blaabjerg, L.I.; Bulduk, B.; Lindenberg, E.; Löbmann, K.; Rades, T.; Grohganz, H. Influence of Glass Forming Ability on the Physical Stability of Supersaturated Amorphous Solid Dispersions. J. Pharm. Sci. 2019, 108, 2561–2569, doi:10.1016/j.xphs.2019.02.028.

23. Blaabjerg, L.I.; Grohganz, H.; Lindenberg, E.; Löbmann, K.; Müllertz, A.; Rades, T. The Influence of Polymers on the Supersaturation Potential of Poor and Good Glass Formers. Pharmaceutics 2018, 10, doi:10.3390/pharmaceutics10040164.

24. Blaabjerg, L.I.; Lindenberg, E.; Löbmann, K.; Grohganz, H.; Rades, T. Is There a Correlation between the Glass Forming Ability of a Drug and Its Supersaturation Propensity? Int. J. Pharm. 2018, 538, 243–249, doi:10.1016/j.ijpharm.2018.01.013.

25. Meng, F.; Ferreira, R.; Zhang, F. Effect of Surfactant Level on Properties of Celecoxib Amorphous Solid Dispersions. J. Drug Deliv. Sci. Technol. 2019, 49, 301–307, doi:10.1016/j.jddst.2018.11.026.

26. Baghel, S.; Cathcart, H.; O’Reilly, N.J. Understanding the Generation and Maintenance of Supersaturation during the Dissolution of Amorphous Solid Dispersions Using Modulated DSC and 1H NMR. Int. J. Pharm. 2018, 536, 414–425, doi:10.1016/j.ijpharm.2017.11.056.

27. Ray, E.; Vaghasiya, K.; Sharma, A.; Shukla, R.; Khan, R.; Kumar, A.; Verma, R.K. Autophagy-Inducing Inhalable Co-Crystal Formulation of Niclosamide-Nicotinamide for Lung Cancer Therapy. AAPS PharmSciTech 2020, 21, 260, doi:10.1208/s12249-020-01803-z.

28. van Tonder, E.C.; Mahlatji, M.D.; Malan, S.F.; Liebenberg, W.; Caira, M.R.; Song, M.; de Villiers, M.M. Preparation and Physicochemical Characterization of 5 Niclosamide Solvates and 1 Hemisolvate. AAPS PharmSciTech 2004, 5, E12, doi:10.1208/pt050112.

29. Worku, Z.A.; Aarts, J.; Singh, A.; Van den Mooter, G. Drug–Polymer Miscibility across a Spray Dryer: A Case Study of Naproxen and Miconazole Solid Dispersions. Mol. Pharm. 2014, 11, 1094–1101, doi:10.1021/mp4003943.

30. Lodagekar, A.; Borkar, R.M.; Thatikonda, S.; Chavan, R.B.; Naidu, V.G.M.; Shastri, N.R.; Srinivas, R.; Chella, N. Formulation and Evaluation of Cyclodextrin Complexes for Improved Anticancer Activity of Repurposed Drug: Niclosamide. Carbohydr. Polym. 2019, 212, 252–259, doi:10.1016/j.carbpol.2019.02.041.

31. Song, Y.; Yang, X.; Chen, X.; Nie, H.; Byrn, S.; Lubach, J.W. Investigation of Drug–Excipient Interactions in Lapatinib Amorphous Solid Dispersions Using Solid-State NMR Spectroscopy. Mol. Pharm. 2015, 12, 857–866, doi:10.1021/mp500692a.

32. Egami, K.; Higashi, K.; Yamamoto, K.; Moribe, K. Crystallization of Probucol in Nanoparticles Revealed by AFM Analysis in Aqueous Solution. Mol. Pharm. 2015, 12, 2972–2980, doi:10.1021/acs.molpharmaceut.5b00236.

33. Que, C.; Lou, X.; Zemlyanov, D.Y.; Mo, H.; Indulkar, A.S.; Gao, Y.; Zhang, G.G.Z.; Taylor, L.S. Insights into the Dissolution Behavior of Ledipasvir–Copovidone Amorphous Solid Dispersions: Role of Drug Loading and Intermolecular Interactions. Mol. Pharm. 2019, 16, 5054–5067, doi:10.1021/acs.molpharmaceut.9b01025.

34. Ueda, K.; Taylor, L.S. Polymer Type Impacts Amorphous Solubility and Drug-Rich Phase Colloidal Stability: A Mechanistic Study Using Nuclear Magnetic Resonance Spectroscopy. Mol. Pharm. 2020, 17, 1352–1362, doi:10.1021/acs.molpharmaceut.0c00061.

35. Kawakami, K.; Sato, K.; Fukushima, M.; Miyazaki, A.; Yamamura, Y.; Sakuma, S. Phase Separation of Supersaturated Solution Created from Amorphous Solid Dispersions: Relevance to Oral Absorption. Eur. J. Pharm. Biopharm. 2018, 132, 146–156, doi:10.1016/j.ejpb.2018.09.014.

36. McConnell, E.L.; Basit, A.W.; Murdan, S. Measurements of Rat and Mouse Gastrointestinal PH, Fluid and Lymphoid Tissue, and Implications for in-Vivo Experiments. J. Pharm. Pharmacol. 2008, 60, 63–70, doi:10.1211/jpp.60.1.0008.

37. Palmelund, H.; Madsen, C.M.; Plum, J.; Müllertz, A.; Rades, T. Studying the Propensity of Compounds to Supersaturate: A Practical and Broadly Applicable Approach. J. Pharm. Sci. 2016, 105, 3021–3029, doi:10.1016/j.xphs.2016.06.016.

38. Wang, X.; Liu, Y.; Shen, C.; Shen, B.; Zhong, R.; Yuan, H. Effect of Particle Size on in Vitro and in Vivo Behavior of Astilbin Nanosuspensions. J. Drug Deliv. Sci. Technol. 2019, 52, 778–783, doi:10.1016/j.jddst.2019.05.005.

39. Wei, Y.; Zhang, L.; Wang, N.; Shen, P.; Dou, H.; Ma, K.; Gao, Y.; Zhang, J.; Qian, S. Mechanistic Study on Complexation-Induced Spring and Hover Dissolution Behavior of Ibuprofen-Nicotinamide Cocrystal. Cryst. Growth Des. 2018, 18, 7343–7355, doi:10.1021/acs.cgd.8b00978.

40. Harmon, P.; Galipeau, K.; Xu, W.; Brown, C.; Wuelfing, W.P. Mechanism of Dissolution-Induced Nanoparticle Formation from a Copovidone-Based Amorphous Solid Dispersion. Mol. Pharm. 2016, 13, 1467–1481, doi:10.1021/acs.molpharmaceut.5b00863.

41. Stewart, A.M.; Grass, M.E.; Brodeur, T.J.; Goodwin, A.K.; Morgen, M.M.; Friesen, D.T.; Vodak, D.T. Impact of Drug-Rich Colloids of Itraconazole and HPMCAS on Membrane Flux in Vitro and Oral Bioavailability in Rats. Mol. Pharm. 2017, 14, 2437–2449, doi:10.1021/acs.molpharmaceut.7b00338.

42. Ricarte, R.G.; Van Zee, N.J.; Li, Z.; Johnson, L.M.; Lodge, T.P.; Hillmyer, M.A. Recent Advances in Understanding the Micro- and Nanoscale Phenomena of Amorphous Solid Dispersions. Mol. Pharm. 2019, 16, 4089–4103, doi:10.1021/acs.molpharmaceut.9b00601.

43. Denninger, A.; Westedt, U.; Rosenberg, J.; Wagner, K.G. A Rational Design of a Biphasic Dissolution Setup—Modelling of Biorelevant Kinetics for a Ritonavir Hot-Melt Extruded Amorphous Solid Dispersion. Pharmaceutics 2020, 12, 237, doi:10.3390/pharmaceutics12030237.

44. Tho, I.; Liepold, B.; Rosenberg, J.; Maegerlein, M.; Brandl, M.; Fricker, G. Formation of Nano/Micro-Dispersions with Improved Dissolution Properties upon Dispersion of Ritonavir Melt Extrudate in Aqueous Media. Eur. J. Pharm. Sci. 2010, 40, 25–32, doi:10.1016/j.ejps.2010.02.003.

45. Klumpp, L.; Leigh, M.; Dressman, J. Dissolution Behavior of Various Drugs in Different FaSSIF Versions. Eur. J. Pharm. Sci. 2020, 142, 105138, doi:10.1016/j.ejps.2019.105138.

46. Sarnes, A.; Kovalainen, M.; Häkkinen, M.R.; Laaksonen, T.; Laru, J.; Kiesvaara, J.; Ilkka, J.; Oksala, O.; Rönkkö, S.; Järvinen, K.; et al. Nanocrystal-Based per-Oral Itraconazole Delivery: Superior in Vitro Dissolution Enhancement versus Sporanox® Is Not Realized in in Vivo Drug Absorption. J. Controlled Release 2014, 180, 109–116, doi:10.1016/j.jconrel.2014.02.016.

47. Chang, Y.-W.; Yeh, T.-K.; Lin, K.-T.; Chen, W.-C.; Yao, H.-T.; Lan, S.-J.; Wu, Y.-S.; Hsieh, H.-P.; Chen, C.-M.; Chen, C.-T. Pharmacokinetics of Anti-SARS-CoV Agent Niclosamide and Its Analogs in Rats. J. Food Drug Anal. 2006, 14, 6.

48. Bergström, C.A.S.; Wassvik, C.M.; Johansson, K.; Hubatsch, I. Poorly Soluble Marketed Drugs Display Solvation Limited Solubility. J. Med. Chem. 2007, 50, 5858–5862, doi:10.1021/jm0706416.

49. Pardhi, V.; Chavan, R.B.; Thipparaboina, R.; Thatikonda, S.; Naidu, V.; Shastri, N.R. Preparation, Characterization, and Cytotoxicity Studies of Niclosamide Loaded Mesoporous Drug Delivery Systems. Int. J. Pharm. 2017, 528, 202–214, doi:10.1016/j.ijpharm.2017.06.007.

